# A mathematical framework for statistical decision confidence

**DOI:** 10.1101/017400

**Authors:** Balázs Hangya, Joshua I. Sanders, Adam Kepecs

## Abstract

Decision confidence is a forecast about the probability that a decision will be correct. Confidence can be framed as an objective mathematical quantity the Bayesian posterior probability, providing a formal definition of statistical decision confidence. Here we use this definition as a starting point to develop a normative statistical framework for decision confidence. We analytically prove interrelations between statistical decision confidence and other observable decision measures. Among these is a counterintuitive property of confidence that the lowest average confidence occurs when classifiers err in the presence of the strongest evidence. These results lay the foundations for a mathematically rigorous treatment of decision confidence that can lead to a common framework for understanding confidence across different research domains, from human behavior to neural representations.

## 1 Introduction

Previous theoretical studies have offered a number of different approaches to understand the statistical and algorithmic issues involved in computing and deploying decision confidence. For instance, a signal detection theory framework is often employed for probing decisions under uncertainty, and can provide a strong basis for understanding decision confidence as well (Fleming and Dolan, 2010; Kepecs et al., 2008; Ma, 2010; Maniscalco and Lau, 2012; Ratcliff and Starns, 2009). Sequential sampling models have been used to understand how decisions are reached based on noisy evidence across time (Bogacz et al., 2006). These can be readily extended with a computation of confidence (Drugowitsch et al., 2014; Pleskac and Busemeyer, 2010; Schustek and Moreno-Bote, 2014; Vickers, 1979). Perhaps the most intuitive extension is within the race model framework, where the difference between decision variables for the winning and losing races provides an estimate of confidence (Kepecs et al., 2008; Merkle and Van Zandt, 2006; Moreno-Bote, 2010; Vickers, 1979; Zylberberg et al., 2012). Mechanistically, neural network models based on attractor dynamics have also been used to study how confidence can be computed by neural circuits (Insabato et al., 2010; Rolls et al., 2010).

Such computational models have also helped to interpret experimental studies on the neural basis of decision confidence. However, it remains unclear how one could identify a confidence computation among mixed signals acquired from the brain, or how confidence in non-human animals can be quantified without verbal reports of their subjective feelings. What would a neural or behavioral implementation of confidence look like with respect to other observable measures of a decision? To resolve this quandary, previous studies employed quantitative models that could provide a formal prediction for what a representation of the internal variable of “confidence” would look like in terms of observable and quantifiable parameters. For instance, the orbitofrontal cortex of rats, a region implicated in the prediction of outcomes, was found to carry neural signals related to confidence (Kepecs et al., 2008). This was established by identifying unique signatures of confidence common to signal detection theory and the race model of decision-making. Similarly, signal detection theory predictions have been used to understand correlates of decision confidence in the dorsal pulvinar (Komura et al., 2013) and sequential sampling models in the parietal cortex (Kiani and Shadlen, 2009). Without such computational foundations, it would not be possible to identify and rigorously study representations of confidence in neurons. Beyond a description of how confidence could be computed, signal detection theory has also been used as a starting point for evaluating the metacognitive sensitivity of human confidence reports (Ferrell and McGoey, 1980; Higham and Arnold, 2007; Higham et al., 2009; Kunimoto et al., 2001; Lachman et al., 1979; Nelson, 1984).

Here we approached the well-studied topic of decision confidence from a mathematical statistics perspective. We had two main goals. First, compared to prior studies, we attempted to make as few assumptions as possible about the structure of noise and decision rule or the algorithm used for estimating confidence. Second, we approached the question of confidence from a psychophysical perspective so it may be useful for psychological and neural studies that often use perceptual uncertainty. The premise of our framework is a normative model of confidence which relates confidence to evidence through conditional probability (Kahneman and Tversky, 1972). While this premise is widely accepted, we show that beyond calibration to outcome probabilities, it makes strong predictions about how a measure of confidence should relate to the discriminability of experimentally presented decision evidence.

We began from first principles in statistics by positing that confidence is a probability estimate describing a belief (Cox, 2006). Thus confidence can be related to the available evidence supporting the same belief through a conditional probability. As such, Bayes’ rule provides a way to understand confidence in terms of quantifiable evidence (Ferrell and McGoey, 1980; Griffin and Tversky, 1992). Formally, decision confidence can be defined as a probability estimate that the chosen hypothesis is correct, given the available perceptual evidence – referred to as the percept. The difficulty with this definition of decision confidence is that it uses the percept, a variable internal to the decision maker. Therefore, it is unclear whether predictions are feasible without explicit assumptions about perception, how the internal percept is generated from the external stimulus. Such assumptions are generally used in the signal detection framework to keep the percept variable mathematically tractable. Here we show, however, that it is possible to analytically derive several novel predictions interrelating confidence with choice correctness and evidence discriminability with few or no assumptions about the percept distribution or about the transfer functions between stimulus, percept and choice.

## 2 Results

From a statistical perspective, a decision process can be viewed as a hypothesis test that evaluates the outcome of a choice against a null hypothesis representing its collective alternatives. Statistical decision confidence can then be defined as a Bayesian posterior probability, which quantifies the degree of belief in the correctness of the chosen hypothesis. In this view, both choice and confidence depend on the quality and amount of evidence informing the particular choice. Therefore we mathematically formalized evidence discriminability based on ideas from psychophysics, as a way to measure the quality of evidence presented to a decision maker.

Based on these definitions, we derived four general properties of statistical decision confidence. First, confidence predicts accuracy: the level of confidence predicts the expected fraction of correct choices – as often intuitively posited. Second, confidence increases with the discriminability of presented evidence for correct choices, but counterintuitively, for incorrect choices, confidence decreases with increasing evidence-discriminability. Third, when presented with a zero-discriminability choice (i.e. an equal amount of evidence supporting each hypothesis, implying chance decision accuracy), the mean decision confidence is precisely 0.75. Fourth, while evidence discriminability itself predicts accuracy (a property referred to as the psychometric function), confidence provides further information improving the prediction of accuracy for any given level of discriminability.

### 2.1 Defining statistical decision confidence

To provide the most general statistical model of a decision process, we define all relevant components (stimulus, percept, choice, confidence) as random variables and the functions that link them (perception, decision) as probabilistic functions. This way the theory presented below applies to both stochastic and deterministic decision rules potentially involving multidimensional stimuli and multiple choices. Decisions are based on an internal variable (decision variable or percept, 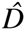), which is the decision makers estimate of a corresponding external variable (stimulus or evidence, *D*).

#### Definition 1.

Let us denote the external variable *D* and realizations of this random variable *d* (referred to as *evidence*). Let us denote the corresponding internal variable 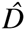 and realizations of this random variable 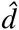 (referred to as *percept*; often referred to elsewhere as the *decision variable*). We define another random variable called the *choice*, denoted by *θ* (realizations denoted by *ϑ*). The choice is a probabilistic function of the percept: 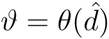.

**Figure 1:**
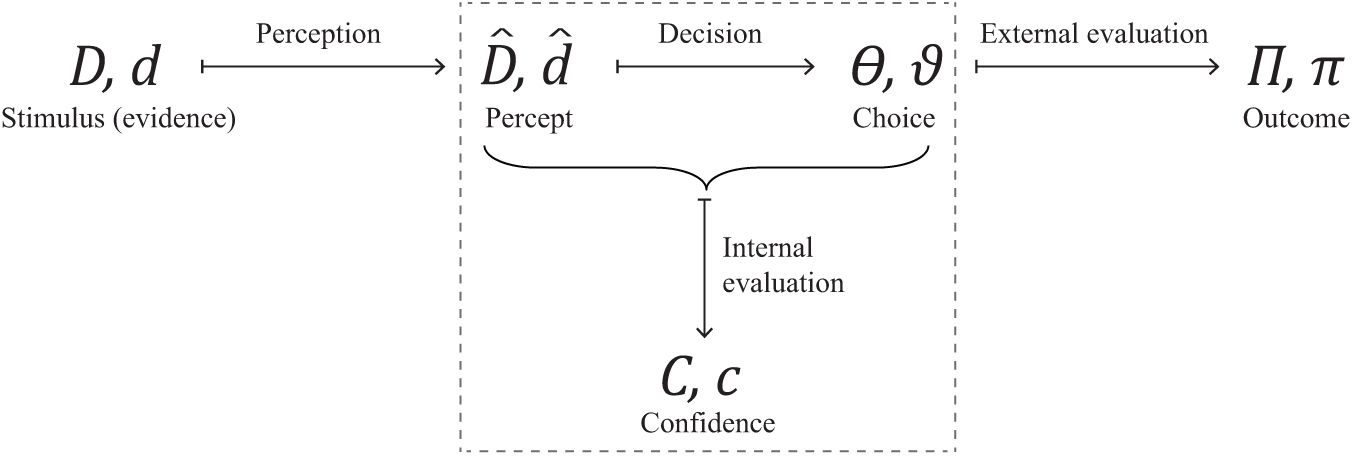
A framework for statistical decision confidence. A stochastic framework of perceptual decision making can be formalized by introducing a small set of random variables. Random variables are denoted by capital letters, and their realizations in lower case.

The choice can be evaluated in terms of a hypothesis testing problem:

null-hypothesis (*H0*): the choice 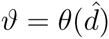 is incorrect;
alternative hypothesis (*H1*): the choice 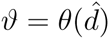 is correct.

Thus, the choice is designated *correct* if the alternative hypothesis is true and *incorrect* otherwise. The evaluation can equivalently be defined as a binary random variable (*outcome*, Π) that is a probabilistic function of choice. Next, confidence (*c*) can be defined as the probability of the alternative hypothesis being true (i.e. Π(*θ*) = 1) provided the percept and the choice.

#### Definition 2.

Define *confidence* as

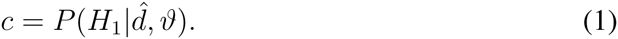

Equivalently,

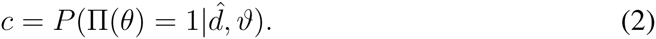

As previously, the random variable will be denoted by *C* and its realizations by *c*. Note that for deterministic choice models, 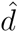 determines *ϑ*, so 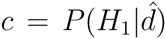. We can now define a function that determines confidence from percept and choice.

#### Definition 3.

Define the *belief function* 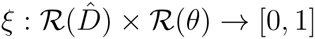 as

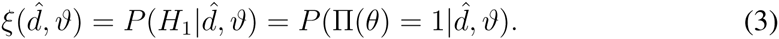

where 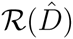 denotes *percept space* and 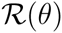 denotes the range of all possible choices (i.e., the *choice space*).

### 2.2 Choice accuracy equals statistical decision confidence

Intuitively, confidence, being defined as an estimate of choice correctness, should predict the expected outcome. We provide a formal treatment of the relationship between confidence and accuracy below.

#### Definition 4.

*Accuracy* is the expected proportion of correct choices:

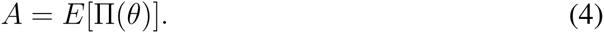

We seek to determine the following function: *f*: [0, 1] → [0, 1], *f*: *c* ↦ *A*c, where *A*c is the accuracy for choices with a given confidence. Our claim is that this function is the identity.

#### Theorem 1.

Accuracy equals confidence:

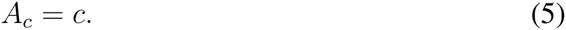

##### Proof.

For every given value of confidence, there is a set of percept-choice pairs leading to the same confidence value: let us denote the image of *c* by the inverse belief function *ξ*^−1^ as 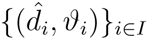, the set of percept-choice pairs mapping onto *c*. Let us first assume that *I* is a countable set. Accuracy for confidence *c* is determined by the probability of a correct choice if *C* = *c* over the probability of encountering the confidence level of *c* (that is, *P* (*C* = *c*)):

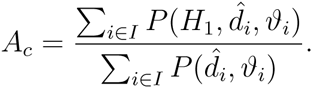

From the definition of joint probability,

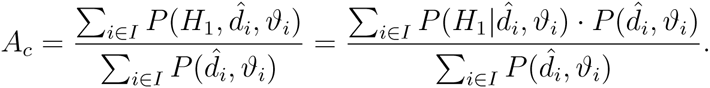

As we know that 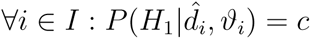,

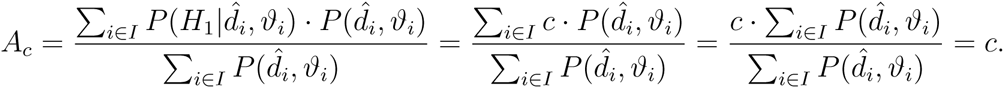

However, *I* is not necessarily a countable set. We can re-write the equations in continuous form to apply to any set as follows.

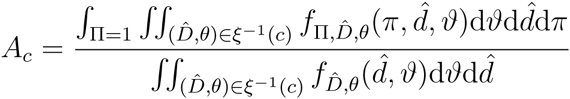

Here Π is a random variable that is 1 if the choice is correct and 0 otherwise (*outcome*, see above).

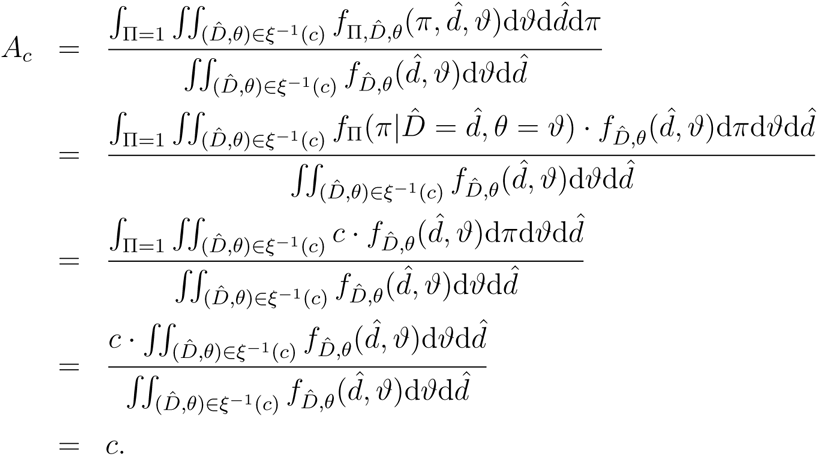

Note that these considerations about confidence do not depend on a particular theory of perception, that is, the function mapping from the external variable (stimulus) onto the internal percept: 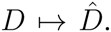 Furthermore, the derivation also does not depend on a particular theory of decision, that is, the function between the percept and the choice: 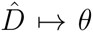. This includes both deterministic and stochastic decision models, the latter referring to models where a percept does not uniquely determine a choice. In case of deterministic decision models, the percept unequivocally determines the choice, thus in the equations we could drop the choice from the inverse picture of confidence, taking only the percept into account: 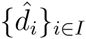 instead of 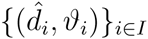. However, as this simplified version would not include stochastic decision models, we chose to adhere to the general formalization.

Another notable aspect of this derivation is that there is no need for a relation to be defined on the percept space. However, if the choice is fixed (or determined by the percept, as in deterministic decision models), confidence defines a natural relation on percepts by *ξ*. More precisely, the order relation on confidence values can be pulled back to the percept space by taking *ξ*^−1^(*c*) and restricting it to a particular choice.

Therefore, we can define the relative terms “low-confidence” percept and “highconfidence percept” based on the relation of confidence values the percepts map onto by the belief function; we will use this concept while proving *Theorem* 2. Please note that this relation always refers to fixed choices.

### 2.3 Confidence increases with increasing evidence discriminability for correct choices and decreases for incorrect choices

Psychophysical studies require to measure decision performance at varying levels of decision difficulty. This necessitates the quantification of the decision difficulty axis, along which the proportion of correct choices can then be measured. Such interrelations, termed psychometric functions, provide a good handle on behavioral performance allowing the detection of subtle changes in behavior. However, there is no single way of grading choice difficulty, resulting in a broad variety of such measures, which complicates the theoretical treatment of psychometric functions. Therefore we define evidence discriminability by its property of measuring difficulty as a class of functions in order to provide a general treatment of the interrelations of choice difficulty and confidence.

#### Definition 5.

Define *evidence discriminability* as a (deterministic) function of the evidence distribution:

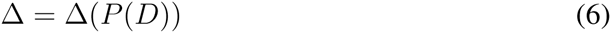

The evidence discriminability function has to fulfill the following property:

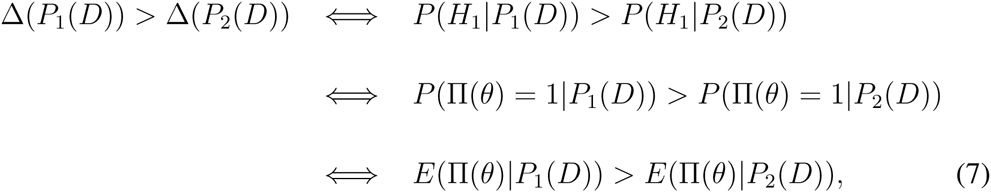

that is, higher discriminability should be equivalent to greater expected outcome (higher probability of correct choices). Any monotonically increasing function of expected outcome satisfies this criterion and can serve as evidence discriminability.

Having defined evidence discriminability, we can now examine how confidence changes with evidence discriminability separately for correct and incorrect choices. We show below that while confidence increases with increasing evidence discriminability for correct choices, it counterintuitively decreases for incorrect choices.

#### Theorem 2.

Let us assume that

- belief independence: the belief function (*ξ*) is independent of evidence discriminability;
- percept monotonicity: for any given confidence *c*, the relative frequency of percepts mapping to *c* by *ξ* changes monotonically with evidence discriminability for any fixed choice.

Under these assumptions, confidence increases for correct choices and decreases for incorrect choices with increasing evidence discriminability.

##### Proof.

We begin with the somewhat counterintuitive claim regarding the incorrect choices.

Let us first examine the two assumptions in more detail.

The first assumption postulates that the function from percept-choice pairs to confidence does not change with evidence discriminability. Thus, whenever we calculate expected value of confidence over a percept distribution, only the percept distributions will depend on evidence discriminability.

For incorrect choices, the second assumption means that with increasing evidence discriminability, the relative frequency of low-confidence percepts increases while the relative frequency of high-confidence percepts decreases in the percept distribution. Note that low-confidence and high-confidence percepts are defined here through the relation imposed by *ξ* on the percepts (see our remark at the end of the previous section). As a trivial consequence of this definition, confidence changes monotonically along low-and high-confidence percepts.

Let us consider two different levels of evidence discriminability (Δ_1_ < Δ_2_), with corresponding distributions of percept restricted to incorrect choices *P* (Δ_1_, low evidence discriminability, i.e. ‘difficult choice’) and *Q* (Δ_2_, high evidence discriminability, i.e. ‘easy choice’). It is sufficient to show that the expected value of confidence is larger for Δ_1_ than for Δ_2_:

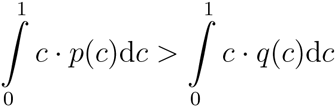

where *p* and *q* denotes the probability density functions corresponding to *P* and *Q*, respectively. Note that *p*(*c*) can be thought of as the probability of the picture of *c* by *ξ*^−1^ restricted to incorrect choices in the percept space.

Equivalently,

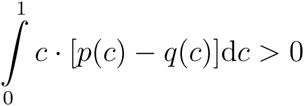

Let *I*_0_ ⊂ [0, 1] denote the interval where *p < q*, and *I*_1_ ⊂ [0, 1] the complementary interval where *p* ≥ *q*. The existence of these intervals is the consequence of the monotonicity assumption. Thus, there is a critical confidence value (denoted here by *c_crit_*) for which *I*_0_ = [0*,c_crit_*] and *I*_1_ = [*c_crit_,* 1]. We then re-write confidence as *c* = *c_crit_* − *c′* if *c < c_crit_* and *c* = *c_crit_* + *c′* if *c > c_crit_*; thus, *c′* > 0 for both cases. Applying these notations,

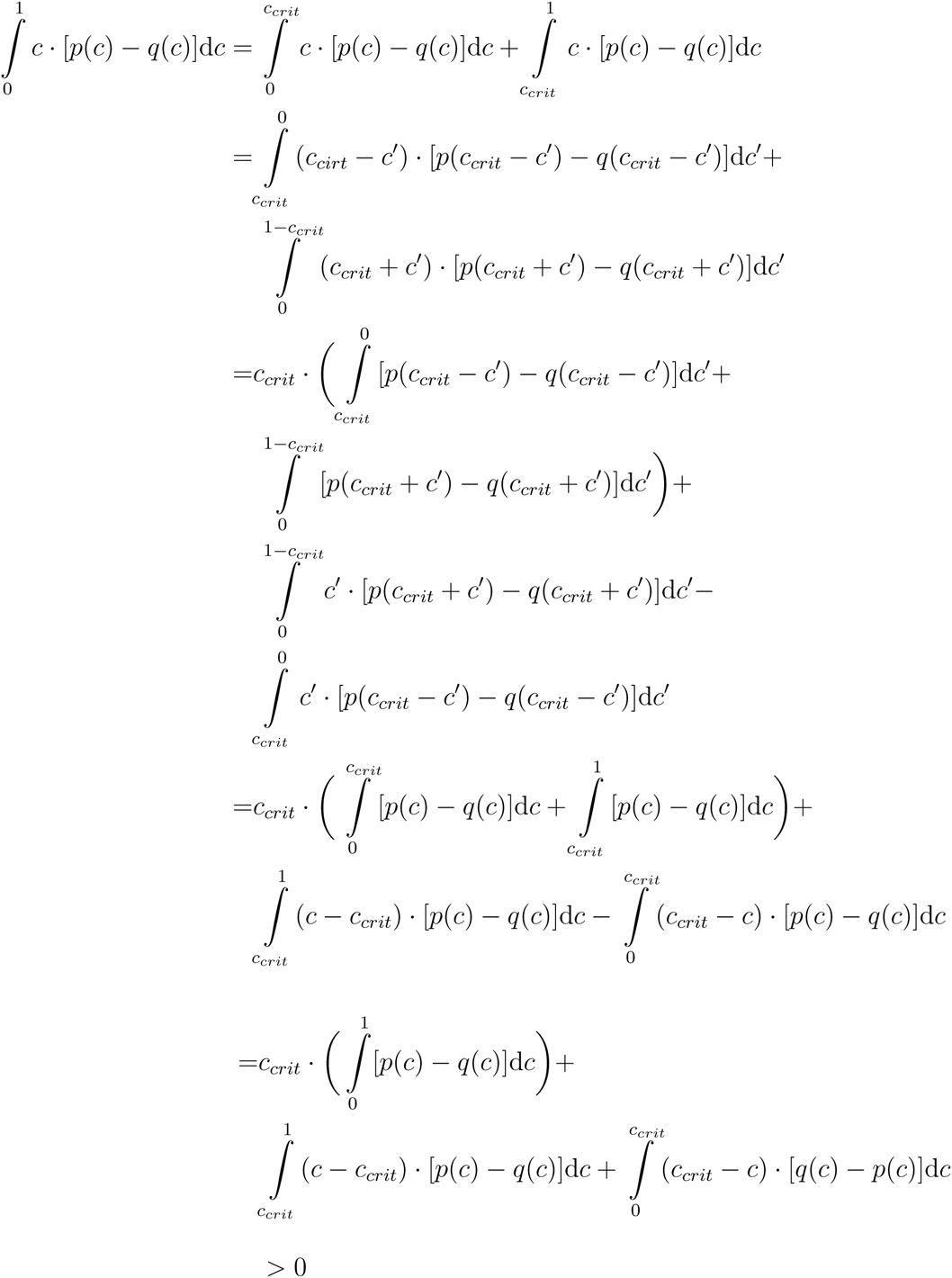

In the last step, the first term is 0, since

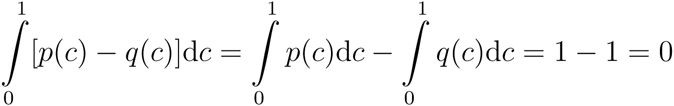

The second term is positive, since *c* − *c_crit_* is positive on *c* ∈ [*c_crit_,* 1] and the probability density functions are evaluated on *I*_1_, where *p* ≥ *q*. Finally, the third term is also positive, because *c*_crit_ − *c* is positive on *c* ∈ [0*,c*_crit_] and the probability density functions are evaluated on *I*_0_, where *q > p*. As a consequence, the sum is positive, which completes the proof for incorrect choices.

For correct choices, high-confidence percepts are increasingly more likely with increasing evidence discriminability, thus present an opposite pattern compared to incorrect choices. Therefore, a symmetric derivation proves the increase of confidence with increasing evidence discriminability for correct choices.

The assumption that *ξ* is independent of evidence discriminability is necessary for this derivation. In this framework, confidence is defined through the true distributions of correct and incorrect choices, kept fixed; therefore this assumption is met. However, if confidence values are updated based on distributions reflecting varying values of evidence discriminability, then the belief function will differ according to evidence discriminability, thus the above proof does not apply. Furthermore, the expected value of confidence cannot decrease with increasing evidence discriminability for incorrect choices: for the lowest levels of discriminability, when the outcome is at chance level, confidence will fall to its lowest possible value, reflecting equal probabilities of the null and alternative hypothesis regardless of the percept. This represents situations in which the decision-maker is provided with information about evidence discriminability, e.g. by grouping decisions of similar discriminabilities (like in a block experimental design), providing an opportunity to learn about evidence discriminability and update the distributions underlying confidence accordingly. Thus, the above theorem only applies when updating confidence based on knowledge of evidence discriminability is prevented, e.g. by randomizing the order of choices with different discriminability levels in an interleaved design.

### 2.4 Confidence predicts outcome beyond evidence discriminability

Psychometric functions reveal accuracy for any given level of evidence discriminability. While confidence also changes with evidence discriminability, it is not obvious whether it carries additional information allowing better prediction of outcome for a given level of evidence discriminability. Below we show that it does.

#### Theorem 3.

For any given evidence discriminability, accuracy for low confidence choices is not larger than that of high confidence choices (splitting the confidence distribution at any particular value). A strict inequality holds in all cases when accuracy is dependent on the percept.

##### Proof.

Let us take the set of low-confidence percept-choice pairs corresponding to the low confidence choices by *ξ*^−1^, and similarly, the set of high-confidence percept-choice pairs corresponding to the high confidence choices. By the definition of confidence (*Definition 2* in *Section 2.1*), low-confidence percept-choice pairs cannot have higher accuracy than the high-confidence percept-choice pairs. If all percepts are associated with the same accuracy (either when the percept does not carry information about the hypotheses of choice, or when the percept determines the correct choice with a probability of one), the two accuracies are equal. Otherwise, the two accuracies should necessarily differ, in which case the strict inequality holds.

Thus, even within the same level of difficulty, the internal noise (e.g. noisy perception) can result in different percepts, some being “easier” and others “harder”. The decision maker has access to this internal variable while the experimenter does not. However, the confidence report contains at least part of this information, providing additional information to the experimenter, which makes the experimenter’s estimate of accuracy better.

### 2.5 The average confidence in neutral evidence

Next, we examine the average confidence at neutral evidence, i.e. evidence carrying no information about the correct choice, for one-dimensional variables.

#### Theorem 4.

Assuming

- the percept is determined by a symmetric distribution centered on the evidence (‘symmetric noise model’),
- the evidence is distributed uniformly over the evidence space, and
- the choice is deterministic,

the average confidence for neutral evidence is precisely 0.75.

##### Proof.

We first prove the following lemma.

#### Lemma 5

Integrating the product of the probability density function and the distribution function of any probability distribution symmetric to zero over the positive half-line results in 3/8:

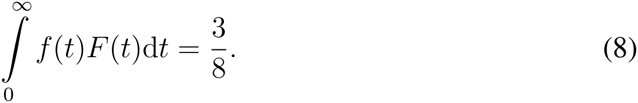

##### Proof.

∀K − ∞ < K < ∞,

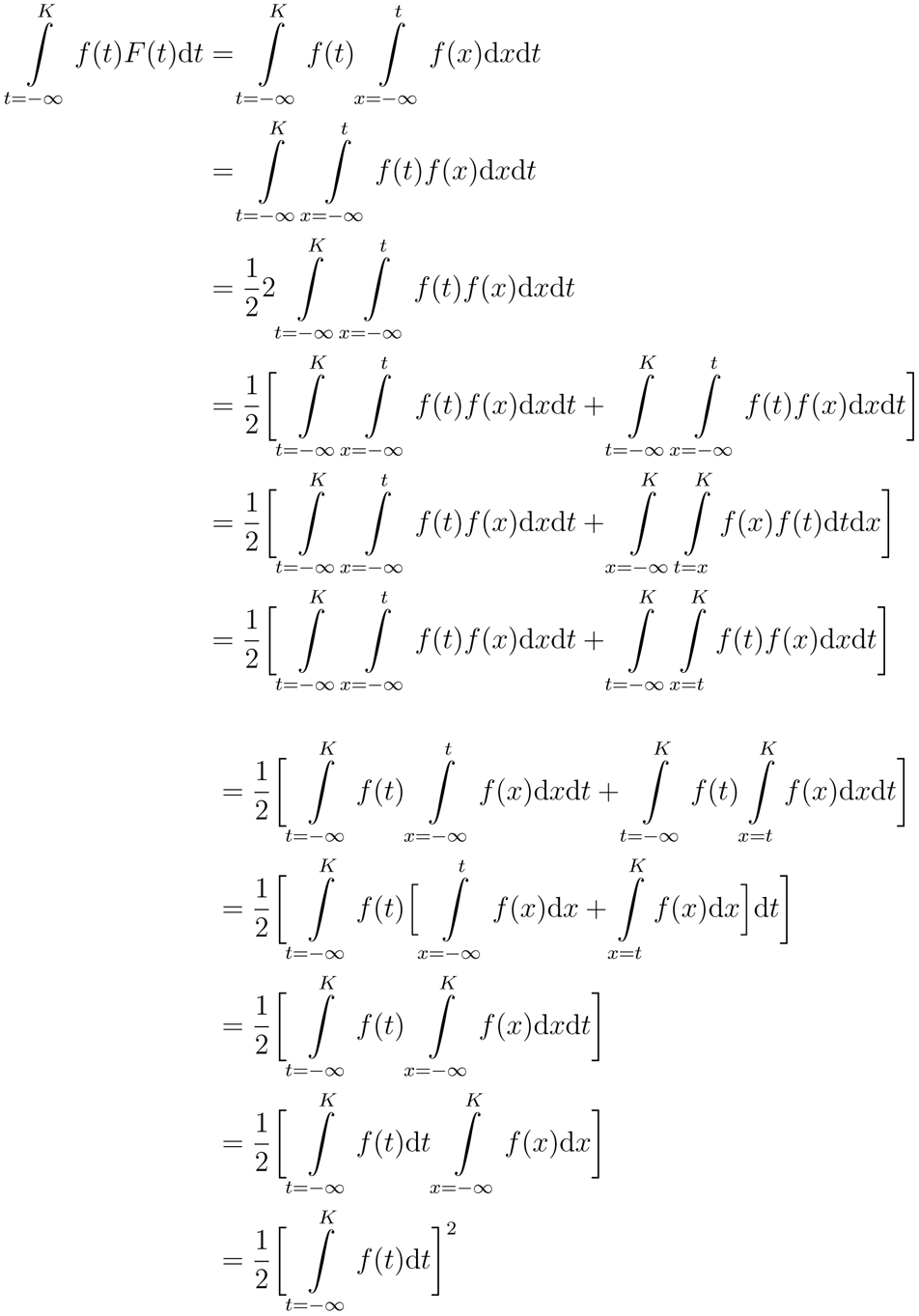

using that
for changing the integral boundaries and then swapping x and t in the second integral term. Applying the above equation, we can write

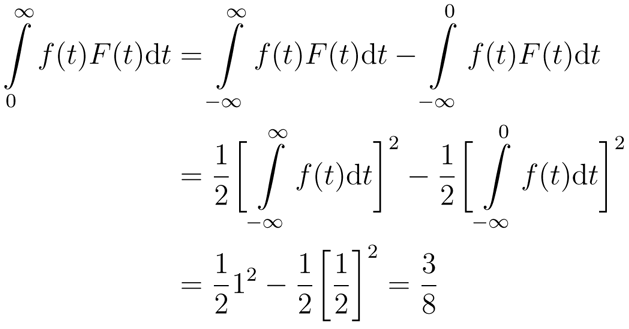

##### Proof of the theorem

Confidence for neutral evidence is determined by the percept corresponding to neutral evidence and the probability of being correct provided the percept and the choice. Thus, the average confidence for neutral evidence can be calculated by integrating over the distribution of percepts provided neutral evidence (indicated here by *d* = 0):

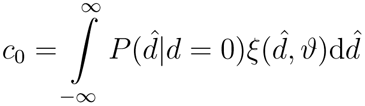

Since we assumed deterministic choice (third assumption), confidence is determined by the percept; therefore we can drop *ϑ* from the equation:

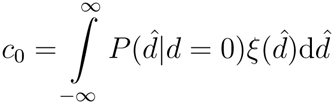

Based on our first assumption, the percept is determined by a symmetric distribution around the evidence. Denote the density function of this symmetric (‘noise’) distribution *f* and its distribution function *F*. Since the percept distribution is symmetric,

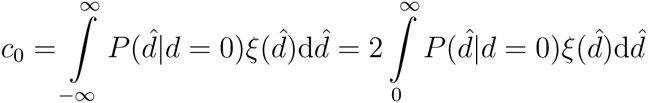

As a consequence of the second assumption of uniform evidence distribution, 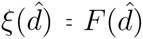 for 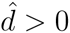:

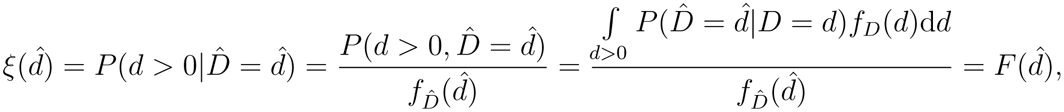

using the theorem of total probability. In the last step, we use that *f_D_* and 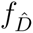 are constant because of the uniformity assumption. (Note that we restrict the support of 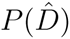 to that of *P* (*D*).) Thus,

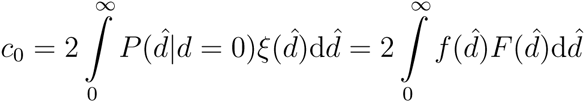

By applying the lemma,

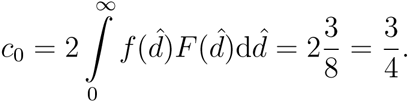

Note that one of the critical assumptions we made is that the evidence is distributed uniformly over the evidence space. While real-life scenarios may often represent non-uniform evidence distributions, this uniformity property holds approximately true for many psychophysics experiments using interleaved evidence strength. Therefore *Theorem 4* provides a quantitatively testable prediction about confidence reports in psychophysics experiments.

### 2.6 Monte Carlo simulations illustrating the signatures of decision confidence

To illustrate our theory we created a Monte Carlo simulation of the normative definition of confidence. For the simulation, we assumed that Gaussian noise (*µ* = 0*, σ* = 0.18) corrupts the external evidence, to generate an internal percept. We used a deterministic decision rule based on the sign of the percept. Thus, outcomes were correct if the sign of the evidence and percept matched. Confidence was calculated as fraction of correct trials for each percept based on *Definition 2*. This enabled us to explore predicted interrelationships between confidence, evidence discriminability and choice. *Figure 2A* shows that confidence predicts the mean choice accuracy (*Theorem 1*). *Figure 2B* demonstrates that mean confidence for a given level of evidence discriminability increases for correct and decreases for incorrect choices (*Theorem 2*), and that the mean confidence for neutral evidence is 0.75 (*Theorem 4*). *Figure 2C* illustrates that for each given level of evidence discriminability, accuracy for high confidence choices is greater than for low confidence choices (*Theorem 3*). Note that while accuracy across simulation trials with low confidence falls to chance (*Figure 2A*), on the converse, the average confidence for neutral evidence is at mid-range (*Figure 2B*). This seemingly contradictory result is explained by the fact that neutral evidence results in a mixture of percepts, most of which are associated with an above-chance average accuracy, leading to an apparent “overconfidence”. Taken together these plots illustrate four signatures of decision confidence in terms of externally quantifiable variables that can be experimentally examined.

**Figure 2:**
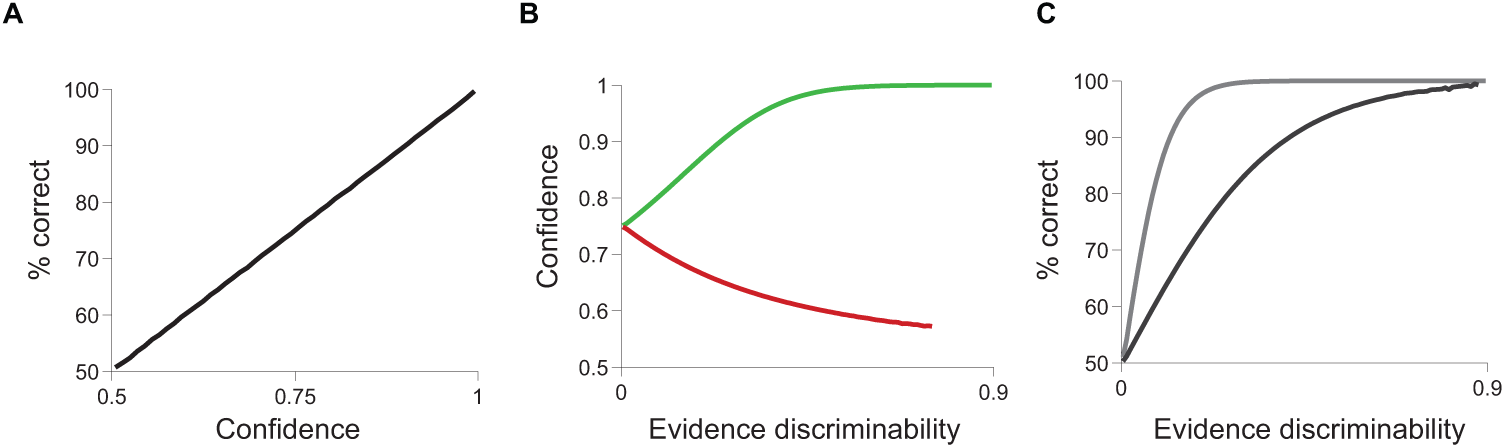
The normative model of confidence predicts specific interrelationships between evidence, outcome and confidence. **(A-C)** Monte Carlo simulations of the normative model (10 billion trials). Bins with fewer than 100 simulation data points were omitted. **(A)** Confidence equals accuracy. **(B)** Average confidence increases with evidence discriminability from 0.75 for correct choices and decreases for errors. **(C)** Conditioning on high or low confidence (split at *c* = 0.8) segregates psychometric performance.

### 2.7 Relating P-values to confidence

Above we derived properties of statistical confidence based on the definition of decision confidence as a Bayesian posterior probability. We next sought to demonstrate their generality by testing their validity on confidence values produced by other statistical approaches. First, we constructed a simulation to test the properties of p-values produced by a common statistical test for evaluating a choice between two hypotheses. First, we examined the one-sided, two-sample Students t-test (*Figure 3A-C*). Samples of 20 measurements were drawn from two Gaussian distributions on each simulation trial, where the simulated task was to identify which underlying distribution had a larger mean. To create graded discriminability, we varied the distance between the means from -0.5 to 0.5 with uniform probability. A simulation trial was designated as “correct” if the mean of the 20 samples drawn from the distribution with the higher mean was higher than the mean of the 20 samples drawn from the distribution with the lower mean. We computed the p-value for each trial using a one sided two-sample t-test to provide a measure of statistical confidence (1 − *p*) in the chosen response. Thus each simulation trial yielded an outcome (correct or error) and a measure of statistical confidence. Second, we also performed simulations of a bootstrap test (Efron and Tibshirani, 1993), which does not depend on a Gaussian assumption about the underlying distributions (*Figure 3D-F*). Exponential sample distributions were used. Offsets for the population means were uniform, ranging between 0 and 1, and the bootstrap sample size was 1000. As shown in *Figure 3*, the p-values derived from a t-test and a two-sample bootstrap test for difference between means reveal the same pattern of interrelationships we derived from the Bayesian confidence definition. Thus, the predictions we derived for statistical decision confidence are valid across different statistical approaches: Bayesian, frequentist and bootstrap statistics.

**Figure 3:**
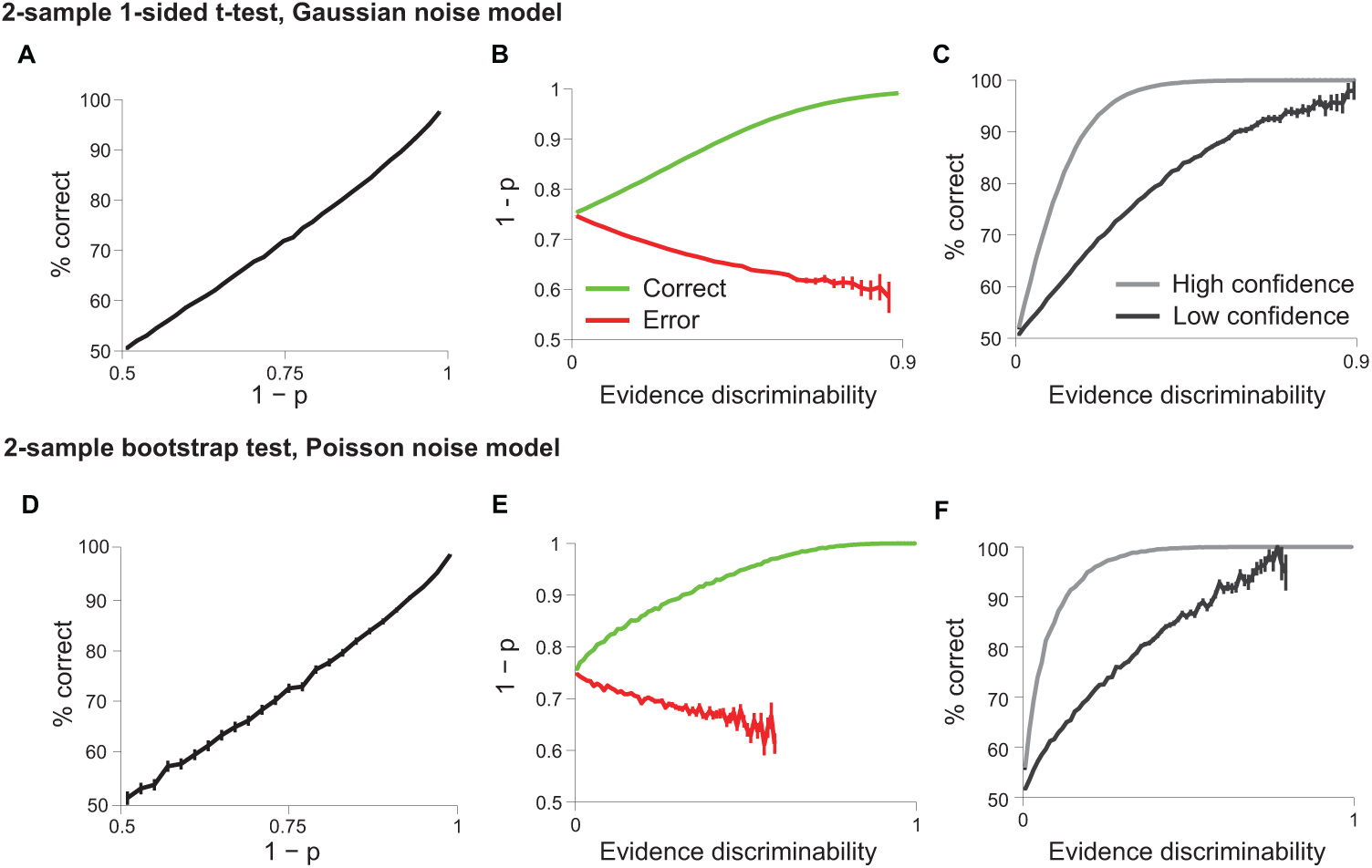
Two statistical tests reproduce patterns predicted by the normative model. **(A-C)** A simulation of 10 million trials evaluated the one-sided, two sample Students t-test p-value with respect to accuracy and evidence discriminability. Since *p* indicates uncertainty, axes show 1 − *p* to indicate confidence. **(A)** 1 − *p* is positively correlated with accuracy. **(B)** 1 − *p* is monotonically increasing with evidence discriminability for correct trials and decreasing for error trials. **(C)** P-values contain information about outcome even at fixed evidence discriminability. **(D-F)** A simulation of the p-value in a one-sided bootstrap test for an ordinal relationship between two means, using exponential distributions.

## 3 Discussion

We presented a normative statistical framework that enables comparisons of statistical decision confidence with confidence measures in other domains. Unlike signal detection theory and other algorithmic frameworks that simulate confidence judgments based on assumptions about the underlying evidence distributions, we show that a strict analytical treatment is possible in a distribution-free manner.

We analytically derived a set of properties of confidence defined as the Bayesian posterior probability of a chosen hypothesis being correct. First, confidence predicts accuracy: the level of confidence predicts the expected fraction of correct choices. This property corresponds most directly to the intuitive notion of confidence as a graded forecast about accuracy. Second, mean confidence for a given level of external evidence is larger for correct than incorrect choices and in fact varies with an opposite sign with evidence discriminability for correct vs. incorrect choices. Specifically, mean confidence levels increase with the ease of discriminability for correct choices, but counterintuitively, confidence decreases with increasing evidence-discriminability for incorrect choices. This surprising dissociation is a consequence of the differences in the distributions of conditional percepts between correct and incorrect choices. Third, and perhaps most surprisingly, when presented with an equal amount of evidence supporting each hypothesis, in other words a non-discriminable choice that will lead to chance accuracy, the mean decision confidence is much greater than chance – precisely 0.75. Fourth, while the psychometric function defines the average choice accuracy for a given level of external evidence, knowledge of confidence provides further information improving the prediction of accuracy for any given level of discriminability.

These four properties are useful for interpreting both behavioral and physiological experiments on decision confidence. Behaviorally, our framework makes it clear that even statistically optimal confidence reports can appear to show systematic miscalibration. This mismatch between confidence reports and accuracy is most dramatically illustrated by the 0.75 mean confidence for neutral evidence that produces chance accuracy behaviorally (0.5). As our framework makes it clear this apparent miscalibration does not imply imperfect prediction of accuracy – rather it is a straightforward consequence of conditioning confidence reports on external variables of the task (e.g,. stimulus difficulty) that are not available to the decision maker. This property of statistical confidence carries important implications for the interpretation of studies demonstrating overconfidence in low discriminability and under-confidence in high discriminability conditions – a controversial phenomenon termed the “hard-easy effect” (Drugowitsch et al., 2014; Ferrell, 1995; Harvey, 1997; Juslin et al., 2000; Merkle, 2009; Moore and Healy, 2008). More generally one has to be careful when analyzing behavior or neural activity by conditioning on external variables not available to the decision maker rather than the internal representations that they are based on. When internal representations are examined as a function of external variables a computational theory is needed to understand how observables conditioned on the external variables is linked to the internal representations. Therefore, rather than revealing miscalibration, conditioning on external variables can be used to test signatures of decision confidence we derived (*Figure 2*) and will be valuable in interpreting putative confidence-related neural activity as well (Kepecs et al., 2008; Komura et al., 2013; Kiani and Shadlen, 2009).

The framework we presented can be interpreted as a prescriptive model, describing how the computation of confidence ought to be done. In this sense it is useful for describing what a neural representation of confidence or its behavioral report should look like. Beyond this, we expect that our mathematical framework will serve as a departure point for quantitatively studying the contribution of confidence to different behaviors, and identifying confidence variables in other domains.

